# Axenic Biofilm Formation and Aggregation by *Synechocystis* PCC 6803 is Induced by Changes in Nutrient Concentration, and Requires Cell Surface Structures

**DOI:** 10.1101/414151

**Authors:** Rey Allen, Bruce E. Rittmann, Roy Curtiss

## Abstract

Phototrophic biofilms are key to nutrient cycling in natural environments and bioremediation technologies, but few studies describe biofilm formation by pure (axenic) cultures of a phototrophic microbe. The cyanobacterium *Synechocystis* sp. PCC 6803 (hereafter *Synechocystis*) is a model micro-organism for the study of oxygenic photosynthesis and biofuel production. We report here that wild-type (WT) *Synechocystis* caused extensive biofilm formation in a 2000 liter outdoor non-axenic photobioreactor under conditions attributed to nutrient limitation. We developed a biofilm assay and found that axenic *Synechocystis* forms biofilms of cells and extracellular material, but only when induced by an environmental signal, such as by reducing the concentration of growth medium BG11. Mutants lacking cell surface structures, namely type IV pili and the S-layer, do not form biofilms.

To further characterize the molecular mechanisms of cell-cell binding by *Synechocystis*, we also developed a rapid (8 hour) axenic aggregation assay. Mutants lacking Type IV pili were unable to aggregate, but mutants lacking a homolog to Wza, a protein required for Type 1 exopolysaccharide export in *Escherichia coli*, had a super-binding phenotype. In WT cultures, 1.2x BG11 induced aggregation to the same degree as 0.8x BG11. Overall, our data support that Wza-dependant exopolysaccharide is essential to maintain stable, uniform suspensions of WT *Synechocystis* cells in unmodified growth medium, and this mechanism is counter-acted in a pili-dependent manner under altered BG11 concentrations.

**Importance:** Microbes can exist as suspensions of individual cells in liquids, and also commonly form multicellular communities attached to surfaces. Surface-attached communities, called biofilms, can confer antibiotic resistance to pathogenic bacteria during infections, and establish food webs for global nutrient cycling in the environment. Phototrophic biofilm formation is one of the earliest phenotypes visible in the fossil record, dating back over 3 billion years. Despite the importance and ubiquity of phototrophic biofilms, most of what we know about the molecular mechanisms, genetic regulation, and environmental signals of biofilm formation comes from studies of heterotrophic bacteria. We aim to help bridge this knowledge gap by developing new assays for *Synechocystis*, a phototrophic cyanobacterium used to study oxygenic phototsynthesis and biofuel production. With the aid of these new assays, we contribute to the development of *Synechocystis* as a model organism for the study of axenic phototrophic biofilm formation.

## 1.0 Introduction

*Synechocystis* is model micro-organism for photosynthesis [1, 2] and is frequently chosen for metabolic engineering to enhance biofuel production (reviewed in [3, 4]. Knowledge of the environmental signals and molecular mechanisms of axenic biofilm formation by microbial phototrophs would inform rational engineering of customized cellular adhesion for biofuel applications, as well as addressing knowledge gaps in the ecological roles of phototrophs in establishing mixed-species biofilms in diverse environments. A few recent studies of axenic phototroph biofilm formation have been reported (reviewed below). However, compared to the wealth of axenic heterotrophic biofilm studies, progress for axenic phototrophs has been limited because the available biofilm assays are relatively slow, and model organisms for axenic phototrophic biofilm formation haven’t been established. As a result, axenic biofilm formation by phototrophs remains poorly understood relative to the prominance of phototrophic biofilms in the earliest fossil record [5–7], and their importance in global nutrient cycling [8–11], astrobiology and space exploration [12–14], and biofuel production [15–17].

We hypothesize that microbial phototrophs, specifically *Synechocystis*, are able to form axenic biofilms, and use cell surface structures such as exopolysaccharides (EPS), S-layer, and pili to attach and adhere to surfaces, similar to other biofilm-forming heterotrophic bacteria (reviewed in [18] and below). To test these hypotheses, we surveyed the literature and used BLAST searches to identify *Synechocystis* genes and predicted homologs known to be important for heterotrophic biofilm formation. These were targeted for deletion in *Synechocystis*.

At neutral pH, bacteria have a net negative charge conferred on them by exopolysaccharides (EPS) (described below). A general model of the role of a cell’s surface charge in binding of cells to a substratum has been described with Derjaguin-Landau-Verwey-Overbeek (DLVO) theory (reviewed in [19, 20]). This model predicts that particles (cells) have a more stable (non-sedimenting) colloid suspension with increasing electrostatic charge of the particles. The extended DLVO model (XDLVO) adds cell surface hydrophobicity to predictions of cell interactions. Proteins, lipids, S-layer glycoproteins, membrane vesicles, and lipopolysaccharides have all been shown to increase the hydrophobicity of a cell surface in a variety of bacteria [21–24], which increase the tendency of cells to aggregate in aqueous (hydrophilic) environments.

Many cyanobacteria synthesize exopolysaccharides (EPS) (reviewed in [25, 26]. Below we highlight three known or hypothesized components of *Synechocystis* EPS: Wza-dependent EPS, the released polysaccharide (RPS) colanic acid, and cellulose. *Synechocystis* EPS have been biochemically characterized using acid hydrolysis and chromatography of the resulting sugar monomers [27, 28]. Selection of naturally occurring mutants by anion-exchange chromatography found that the increasing negative charge of mutants correlated with either increased uronic acids or increased total amount of EPS. Additionally, these strains had decreased rate of cell sedimentation, suggesting that the net charge conferred by EPS influences cell-cell repulsion [29] in *Synechocystis*.

Bacterial EPS synthesis and export systems are best characterized in *Eschericia coli* (*E. coli*). This includes Type I capsular polysaccharide, which requires proteins including Wzx, Wzy, Wza, and Wzc for the later stages of EPS synthesis and export [30]. Wza and Wzc proteins act as a gating mechanism / polymerase, and an outer membrane porin of Type I capsule in *E. coli*, respectively [31]. In many heterotrophic bacterial species, mutants lacking Wza do not synthesize capsule [32] and are also impaired for biofilm formation [33–35]. The *Synechocystis* protein Sll1581 is a putative homolog to Wza (28% identity, 42% similarity), consistent with its localization to the outer membrane of *Synechocystis* [36]. In a previous study, *Synechocystis* mutants lacking Sll1581 had EPS levels less than 25% that of wild-type (42). Compared to WT cells, this mutant showed spontaneous auto-sedimentation without aggregation in supernatants, as measured by a 3-week standing flask assay of liquid cultures. A second putative *Synechocystis* homolog, Sll0923, has 21% identity and 39% similarity to *E. coli* protein Wzc. *Synechocystis* mutants lacking the Wzc protein had the same non-sedimenting phenotype as WT in standing flasks of supernatant (42), and correspondingly only 50% reduction in EPS levels compared to WT. We did not find putative homologs to Wzx and Wzy in *Synechocystis*, but predict the activities of these proteins in Wza-dependent EPS may be provided by ABC-transporters, similar to Type 2 and Type 3 capsule in *E. coli*, as suggested previously [28].

Colanic acid, a type of released polysaccharide (RPS), is synthesized and exported using the same proteins as Type I capsule in *E. coli*, but it is under different genetic regulation. Colanic acid is produced by *E. coli* strains that do not synthesize Wzi, a protein specific to anchoring Type 1 capsule to the cell surface [30, 37]. We did not find a homolog to Wzi in *Synechocystis* using BLASTP search. This is consistent with previously reported detection of RPS in WT *Synechocystis* supernatants, and reduced levels of this RPS in supernatants of *wzc* mutant cultures [29, 38]. The role of *Synechocystis* RPS in cell binding, if any, is not known.

Cellulose is a component of EPS associated with the cell surface, promoting adhesion in many heterotrophic bacteria [39]. Cellulose has also been detected in the EPS of several cyanobacterial species [40, 41]. Instead of cellulose synthases like BcsA that are common in heterotrophic bacteria, cyanobacteria use CesA, the cellulose synthase protein conserved in higher plants [42, 43]. *Synechococcus* PCC 7002 [41] and *Thermosynechococcus vulcanus* RKN [40, 44] were shown to have cellulose-dependent aggregation. *Synechocystis* contains one cellulose synthase motif, DDD35QXXRW, in Sll1377, which has homology to the N-terminal region (48% query coverage) of CesA in *Thermosynechococcus vulcanus* RKN (BLASTP search [45] using default parameters). In a study of 12 diverse cyanobacterial species, cellulose was not detected in *Synechocystis* or *Synechococcus elongatus* PCC 7942 (49) in non-aggregated cultures. From these findings, it is inconclusive whether *Synechocystis* synthesizes cellulose during aggregation.

A second line of evidence in the literature ties cellulose-dependent aggregation to environmental signals via highly conserved role of secondary messenger c-di-GMP (cyclic di-guanosine monophosphate) [46, 47]. Nutrient limitation such as carbohydrate starvation is frequently correlated with increased c-di-GMP levels in heterotrophs [48]; carbohydrate starvation results in both nutrient and energy limitation in heterotrophs.

The cyanobacterium *Thermosynechococcus vulcanus* RKN undergoes cellulose-dependent aggregation in response to blue light [40, 44], an energy-limited condition, in a process requiring formation of c-di-GMP by the protein SesA (Tlr0924). Two studies show that c-di-GMP levels are correlated with aggregation in *Synechocystis* under different energy-limiting conditions, although cellulose measurements were not reported [49, 50].

In addition to the role of EPS in biofilm formation and aggregation, we also investigated the roles of S-layer and pili. The *Synechocystis* S-layer protein (Sll1951) is glycosylated and forms a surface layer with hexagonal symmetry, which can be imaged via TEM as a honeycomb-like surface texture on WT cells [51, 52]. Heterotrophic S-layer mutants have a range of adhesion phenotypes, ranging from super-binding to completely biofilm deficient, depending on species (reviewed in [21]). S-layer mutants have enhanced biofilm formation compared to WT (a super-binding phenotype) in *Bacillus cereus* [53], *Caulobacter crescentus* [54], and various *Clostridium difficile* 630 strains [55]. In contrast, S–layer mutants of *Streptococcus gordonii* are deficient in aggregation [56].

There are six major classes of pili (and / or homologous structures called fimbriae and curli), which pathogens such as *E. coli*, *Pseudomonas aeruginosa*, *Salmonella enterica*, and *Neisseria* species use for attachment, adhesion, and biofilm formation during infection (reviewed in [57]). Additionally, biofilm initiation depends on motility in many bacterial species (reviewed in [58], including gliding motility conferred by Type IV pili. *Synechocystis* has thousands of Type IV pili arranged peritrichously and extending several microns beyond the cell surface [29, 59]. These Type IV pili are glycosylated along their entire length. Mutations causing altered glycosylation of PilA, the pilin structural subunit, cause defects in gliding motility in *Synechocystis* [60]. PilC is a predicted cytoplasmic chaperone protein for export of PilA monomer for pilin assembly in diverse bacteria. Consistent with this prediction, *Synechocystis pilC* deletion mutants (*slr0162-0163*) are apiliate (bald) [61] and amotile. Interestingly, knockout of Pcc7942 2069, a putative homolog of Type II secretion protein E and Type IV pili assembly PilB, causes biofilm formation in *Synechococcus elongatus* PCC 7942 [62] (hereafter *S. elongatus*). This mutation is proposed to block an unidentified molecule that inhibits secretion of biofilm enhancing proteins in WT *S. elongatus* [63]. A subsequent study found certain piliated *S. elongatus* mutants also underwent a degree of sedimentation, adhesion, and biofilm formation compared to WT, indicating that loss of Type IV pili is not a prerequisite for biofilm formation in *S. elongatus* [64].

In this study, we document extensive biofilm formation in a large outdoor photobioreactor used to grow WT *Synechocystis*. We adapted the crystal violet assay commonly used for biofilm study of heterotrophic bacteria in order to screen conditions leading to biofilm formation by axenic WT *Synechocystis* cultures. We also developed rapid aggregation and flocculation assays to further characterize environmental signals and cell surface biochemistry of cell binding. We engineered targeted genetic mutations of genes *sll1581* (*wza*), *slr0923* (*wzc*), *sll1951* (S-layer), and *slr0162-0163* (*pilC*) required for cell surface structures. We then screened these mutants for biofilm formation and aggregation phenotypes. Finally, we compared biofilm formation, aggregation, and motility phenotypes of WT and mutant straints with measurements of outer membrane proteins, and cellulose content of extracellular matrices. We summarized our findings and interpret their significance to biotechnology and microbial ecology.

## 2.0 Results and Discussion

### 2.1 Biofouling in Outdoor PBR is Correlated With Use of Hard Water to Prepare BG11

We grew wild-type *Synechocystis* cultures in a 2000-liter outdoor photobioreactor (PBR). As shown in Figure 1, when cultures were inoculated into growth medium BG11 prepared from sanitized (but not axenic) hard (tap) water, we observed consistent and florid growth of macroscopic green biofilms (n = 3; PBR cleaned and sanitized between replicates). These unwanted biofilms (biofouling) occurred in stages characteristic of heterotroph biofilms that began with isolated colonies and progressed to thick confluent growth over approximately one week of mid-log culture growth. Visible biofilm only grew in illuminated areas, suggesting that this process was driven by obligate phototrophs such as *Synechocystis* [65], and not other microbes present in this non-axenic environment. In subsequent rooftop PBR cultures in BG11 prepared with softened tap water, visible biofilm was absent (not shown). Our data are consistent with the hypothesis that *Synechocystis* formed biofilms only when nutrient levels were reduced due to precipitation with calcium in hard water.

**Fig 1.**
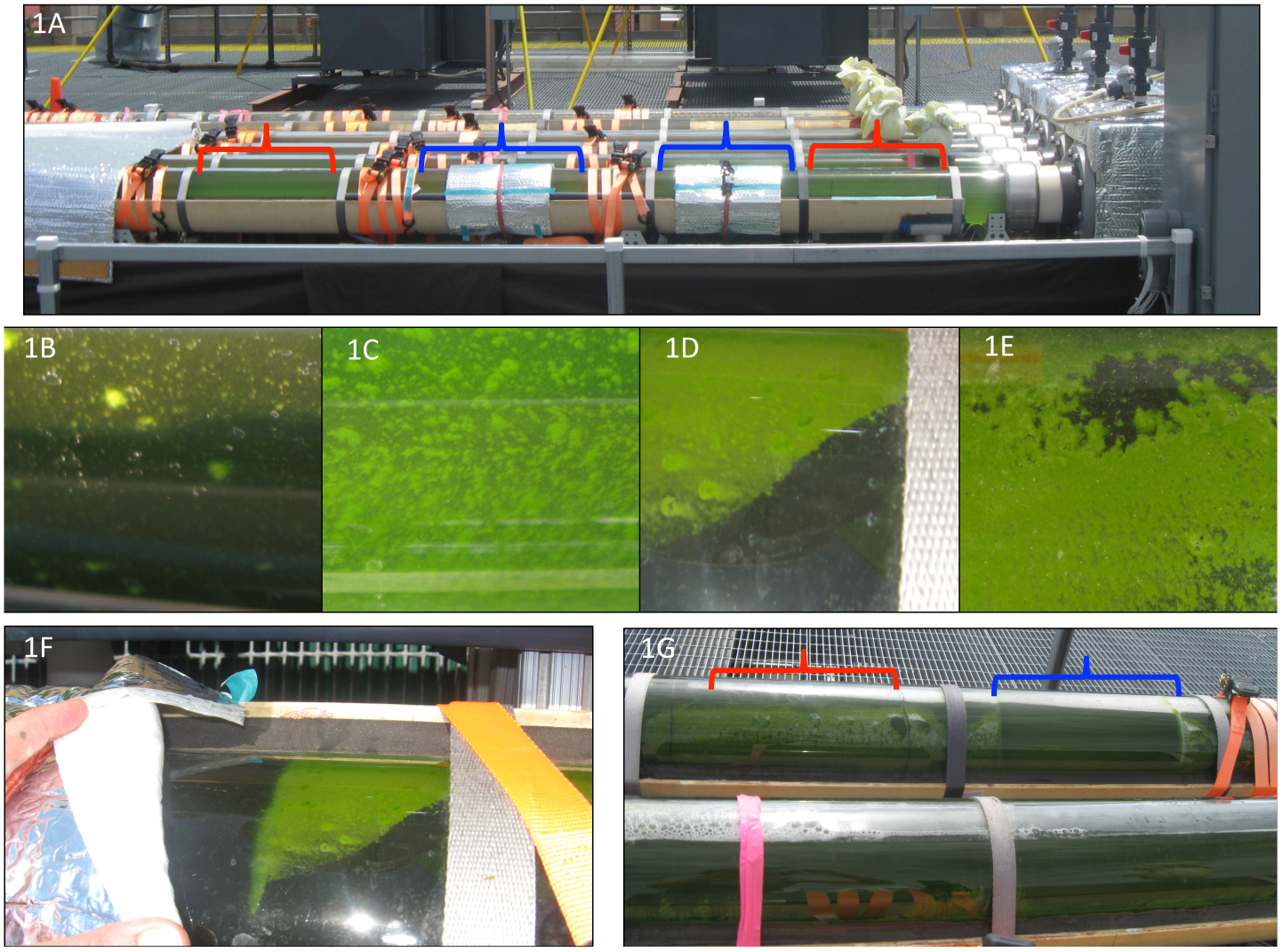
Biofouling of Non-Axenic Roof-Top Photobioreactor During Growth of WT *Synechocystis*. Mylar sheets were used to block light in some regions of the roof–top photobioreactor (RT–PBR) to test the effect of illumination on macroscopic biofouling (unwanted biofilm formation). These covered regions (indicated in blue brackets) were compared with positive control regions that lacked mylar sheets (red brackets). Fig. 1A: During lag phase, no biofilm growth was evident. Fig. 1B to Fig. 1G: Images taken every 24 hours during rapid growth (approximate doubling every 24 h over a period of five days). Biofouling such as representative images in Figure 1 was correlated with using hard tap water to prepare BG11; no biofouling was evident when softened tap water was used.

### 2.2 *Synechocystis* Forms Axenic Biofilms When Concentration of BG11 is Altered

To study axenic *Synechocystis* biofilm formation under controlled laboratory conditions, we adapted the crystal violet assay typically used to study heterotrophic biofilm formation [66, 67] by introducing nutrient evaporation and dilution steps that induce *Synechocystis* biofilm formation. We used this modified crystal violet assay to screen *Synechocystis* biofilm formation in a range of growth conditions including 0-120 rpm shaking, 4-50 μmol/(m^2^ s) photons (PAR, photosynthetically active radiation), with biofilm growth measured daily for up to 5 days (data not shown). In Figure 2, maximum biofilm formation, as determined by crystal violet staining, occurred under conditions of 32 μmol/(m^2^ s) photons (PAR), 30°C, and 72 rpm shaking, at 72 hours (‘Treatment’ condition). It is noteworthy that unlike many heterotrophic bacteria, *Synechocystis* did not form visible biofilms during the crystal violet assay unless induced by these changes in nutrient concentration (p-value = 0.02). Simply diluting cultures as for sub-culturing, without the preceding evaporation step, did not induce biofilm formation (‘Control’ condition; see Materials and Methods section). Scanning laser confocal microscopy in Figure 2B revealed that isolated micro-colonies were approximately 200-300 microns wide, 1-2 cells tall, and uniformly distributed on the submerged portion of the coverslips. Figure 2C shows that stained material bound to coverslips included cells and extracellular material.

**Fig 2.**
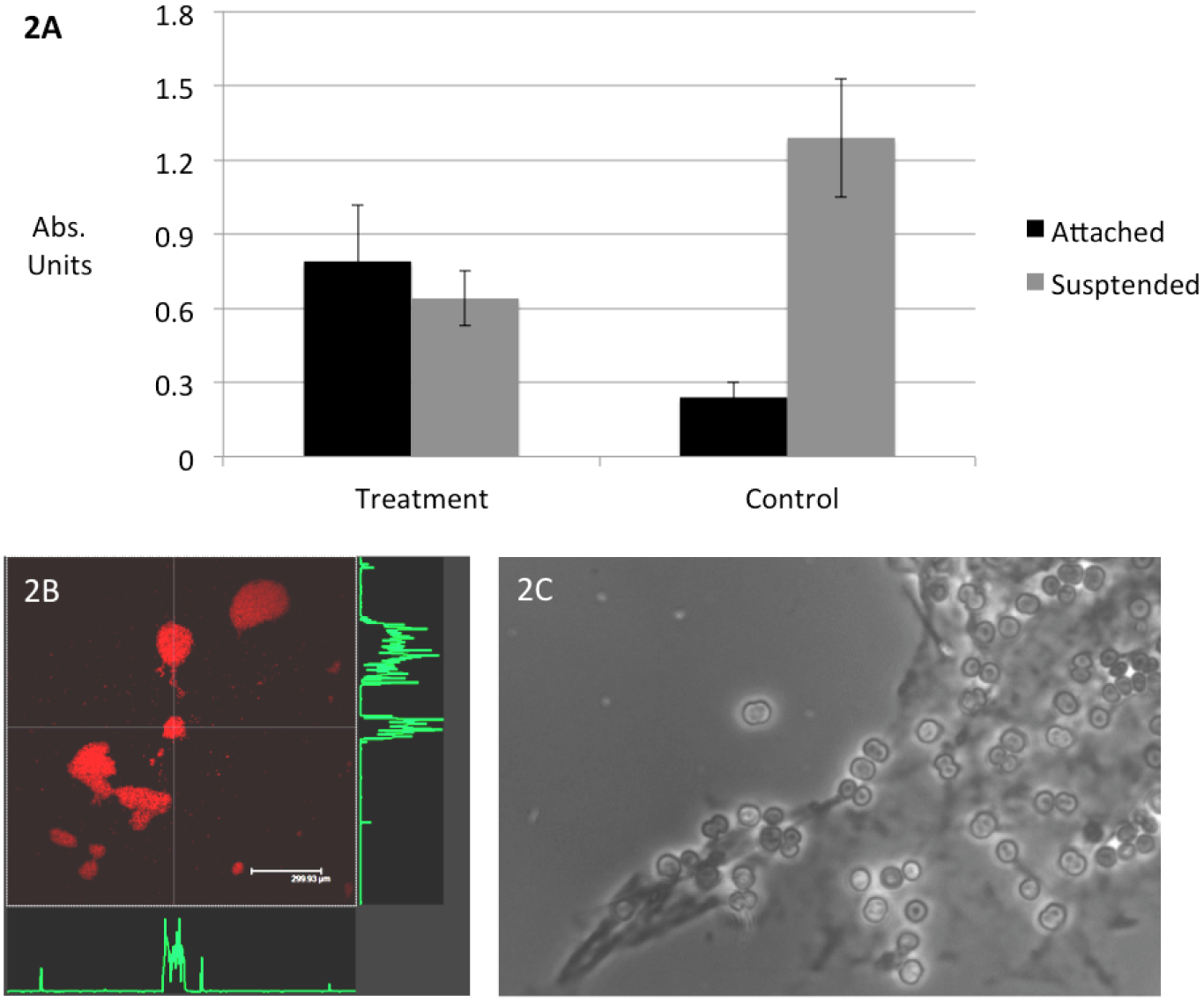
Axenic Biofilm Formation by WT *Synechocystis* Requires Shift in Nutrient Concentrations. Fig. 2A: ‘Attached’ data series show axenic biofilm formation grown from 3 mL cultures on glass coverslips in 12-well tissue culture plates. Standard crystal violet assay had only background (media-only) levels of crystal violet staining (‘Control’ condition), as measured by crystal violet absorbance at OD_600_. Modifying the assay to include evaporation followed by dilution of media induced biofilm formation (‘Treatment’). Crystal violet was eluted from cellular material bound to glass coverslips after 72 h of biofilm growth at 32 μmol/(m^2^ s) photons at 72 rpm shaking. Planktonic cells (‘Suspended’ data series) were also measured at 72 hours at OD_730_. Error bar correspond to one standard deviation from sample mean. Fig. 2B: Scanning laser confocal microscopy of auto-fluorescent cells showing biofilm structure (Bar = 300 microns) approximately 1-2 cells tall, as measured by calibrated Z-axis and indicted with green trace lines (not to scale) showing height of microcolonies at white cross-sections. Fig. 2C: Phase contrast microscopy showed material stained with crystal violet corresponded with attached cells and extracellular material (1000x magnification).

### 2.3 *Synechocystis* Requires Type IV Pili and S-layer to Form Biofilms

We used allelic exchange of the KmR*sacB* markers with genes essential for Type IV pili (*pilC*, (*slr0162-0163*)), EPS (*sll1581*, *sll0923*), and S–layer (*sll1951*) to engineer mutant strains. Three independent isolates of each fully segregated clone were confirmed by PCR and sequencing. Strains and plasmids used in this study are listed in Table 1; primers are listed in Supplementary Material Table S1. We assessed the biofilm phenotypes of mutant strains using the modified crystal violet assay described above. For each condition, we measured four biological replicates (unique cultures). At least four biofilm coupons served as technical replicates for each biological replicate. Figure 3 shows that crystal violet staining from the *pilC* mutants (SD519) (OD_600_ of 0.23±0.02, p-value = 0.02) and S-layer mutants (SD523) (OD_600_ of 0.11±0.03, p-value = 0.01) were significantly lower than WT (SD100) (OD_600_ of 0.78±0.23). We conclude that Type IV pili and S-layer are essential for biofilm formation by *Synechocystis*. Our Wza deletion mutants (SD517) appeared to have a growth defect (OD_730_ of 0.39±0.05) compared to WT (OD_730_ of 0.64±0.11, p-value = 0.01). Growth and/or energy is required for biofilm formation in certain other bacteria [58, 67]. We could not determine whether growth or Wza-dependent EPS is required for biofilm formation by *Synechocystis* from these data.

**Table 1.** Strains and Plasmids Used in this Study.

| Strain | Genotype | Note | Source |
| --- | --- | --- | --- |
| SD100 | WT Kazusa | wild-type, amotile [68] | Gift from Willem Vermaas Lab |
| SD500 | WT Paris | wild-type, motile [68] | Pasteur Cyanobacterial Collection |
| SD515 | Paris sll1581::KmR*sacB* | *wza* mutant | This study |
| SD517 | Kazusa sll1581::KmR*sacB* | *wza* mutant | This study |
| SD519 | Kazusa slr0162-0163::KmR*sacB* | *pilC* mutant | This study |
| SD520 | Paris slr0162-0163::KmR*sacB* | *pilC* mutant | This study |
| SD522 | Paris sll1951::KmR*sacB* | S-layer mutant | This study |
| SD523 | Kazusa sll1951::KmR*sacB* | S-layer mutant | This study |
| SD546 | Kazusa sll0923::KmR*sacB* | *wzc* mutant | This study |
| SD577 | Kazusa slr0162-0163::KmR*sacB* + pΨ552 | *pilC* mutant with extra-chromosomal *pilC* | This study |
| Plasmid | Genotype | Note | Source |
| pGEM 3Z | pUC18 derivative | cloning vector | Promega Corporation |
| pΨ540 | *sll1581* up::KmR*sacB*::*sll1581* down | suicide vector for constructing SD515, SD517 | This study |
| pΨ541 | *slr0162-0163* up::KmR*sacB*::*slr0162-0163* down | suicide vector for constructing SD519, SD520 | This study |
| pΨ543 | *sll1951* up::KmR*sacB*::*sll1951* down | suicide vector for constructing SD522, SD523 | This study |
| pΨ546 | *sll0923* up::KmR*sacB*::*sll0923* down | suicide vector for constructing SD546 | This study |
| pΨ552 | *slr0162-0163* | expression vector for *pilC* | This study |
| pΨ568 | RSF1010 derivative | expression vector | Gift from Soo-Young Wanda |
| pBSBA2KS | pSL1180 derivative (Pharmacia Biotech) | source of KmR*sacB* markers [69] | Gift of Willem Vermaas lab |

**Fig 3.**
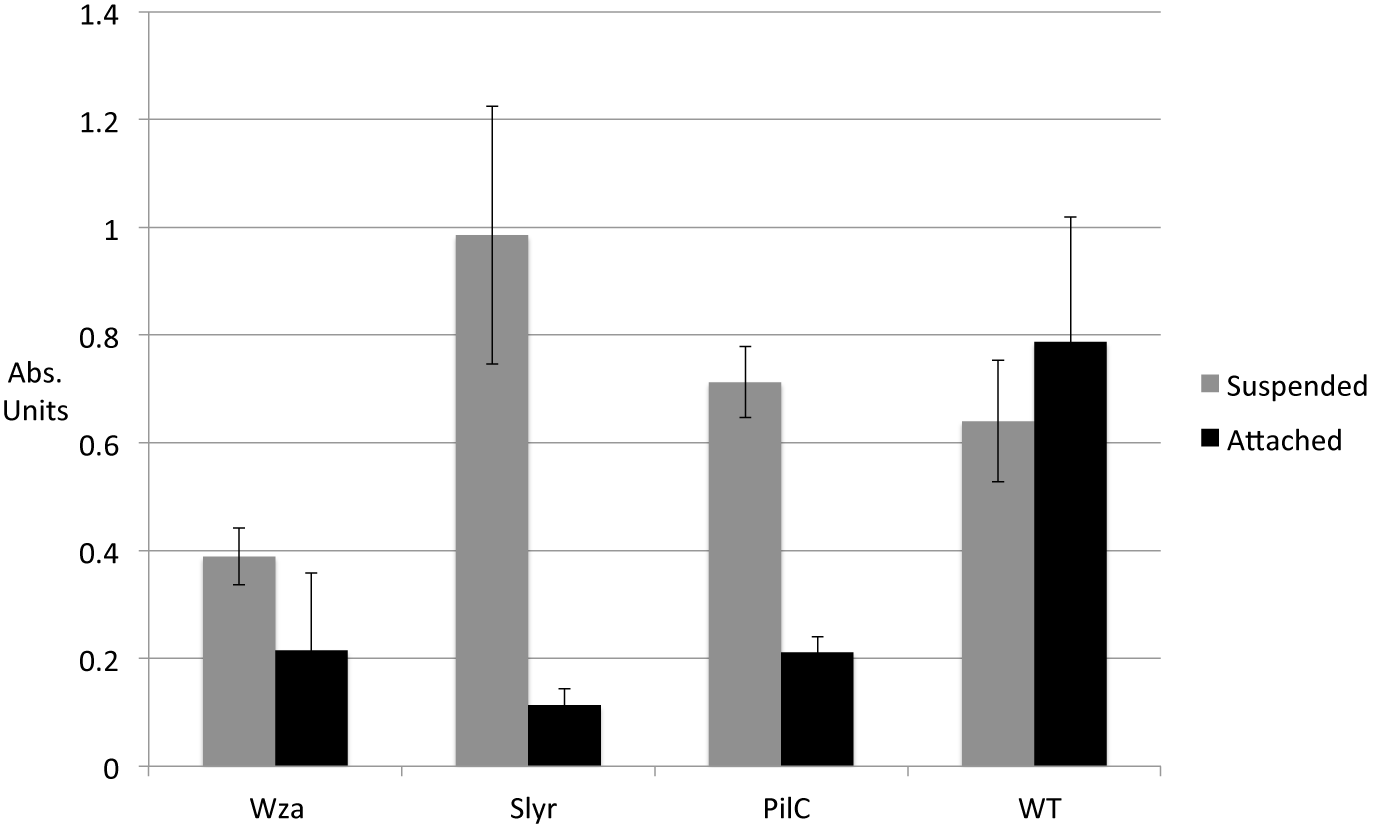
S–layer and Type IV Pili Are Required For Biofilm Formation by *Synechocystis*. ‘Attached’ data series show crystal violet binding measured at OD_600_. ‘Suspended’ data series show planktonic growth measured at OD_730_. ‘Wza’ = SD517; ‘Slyr’ = SD523; ‘PilC’ = SD519; ‘WT’ = SD100. Error bar corresponds to one standard deviation from sample mean.

### 2.4 *Synechocystis* Aggregation Requires Cellular Energy Production

Our crystal violet data show that pili and S-layer are necessary but not sufficient for biofilm formation: presence of these surface structures did not cause WT cells to form biofilms unless some additional unknown factor was induced, such as by changes in nutrient concentrations. We wanted to improve our understanding of the environmental signals and molecular mechanisms of cell-cell binding in *Synechocystis*. Aggregation is related to biofilm formation in that it also involves cell-cell binding, and results in multi-cellular structures. Aggregates and biofilms are both relevant to many of the same ecological processes and biotechnology applications [70, 71]. Additionally, the amount of cellular material in the *Synechocystis* biofilms we grew on coverslips was insufficient for convenient biochemical analyses, likely due to the small culture volume (3 mL per biofilm coupon). Therefore, we developed a rapid aggregation assay to further characterize cell-cell binding by *Synechocystis*.

Figure 4 shows that WT *Synechocystis* cultures with a starting OD_730_ = 0.5 to 0.8. aggregated an average of 56±6 % total biomass within eight hours when shifted to reduced–strength medium (0.8x BG11), compared to negative control cultures resuspended in supernatant (p-value<0.01). This is a similar result to our biofilm data (Fig. 2A). Our data are consistent with reports of a marine cyanobacterium, *Synechococcus* sp. WH8102, producing aggregates when either phosphorus or nitrogen concentrations were lowered [72, 73]. Similarly, lowering iron or phosphorus nutrient concentration induced aggregation and synthesis of extracellular material by the marine cyanobacterium *Trichodesmium* IMS101 [74].

**Fig 4.**
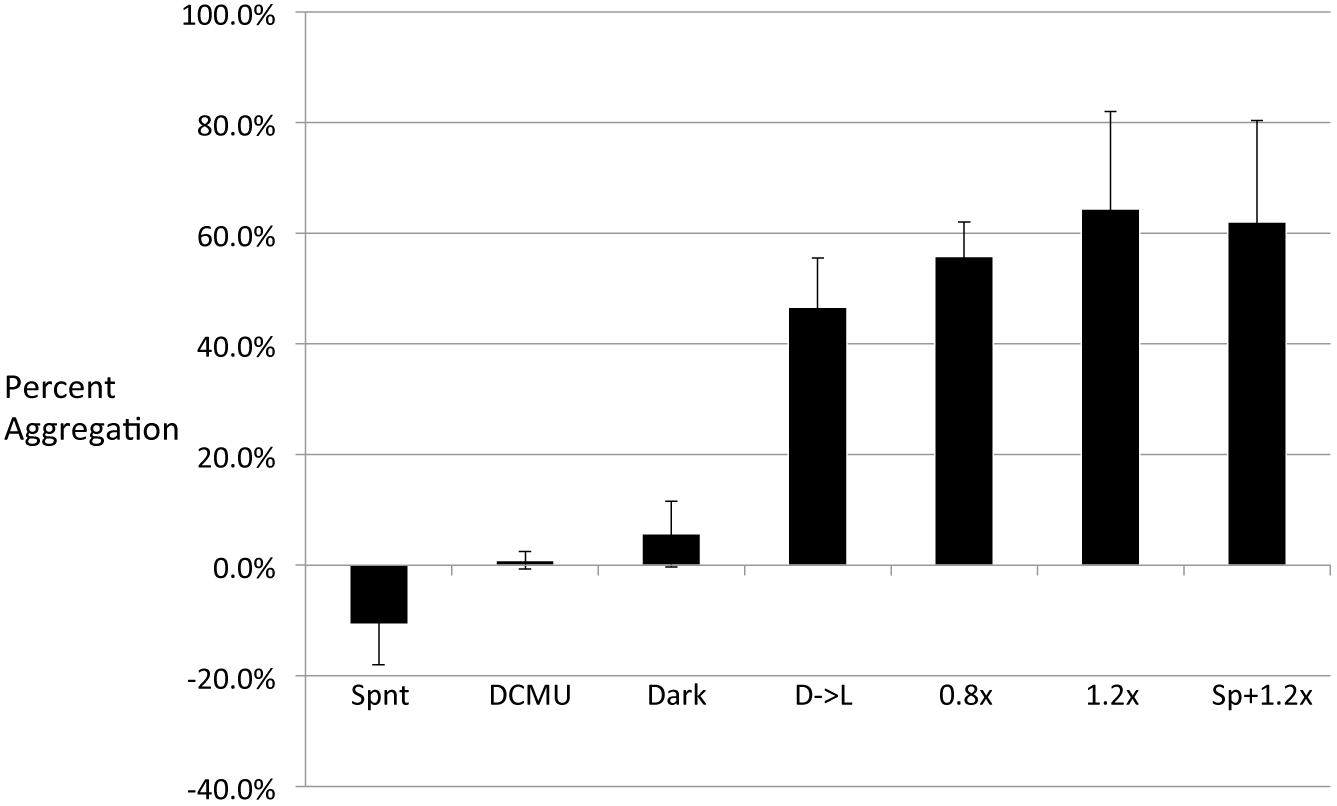
Aggregation of WT *Synechocystis* Requires Cellular Energy Production and is Not Affected by Removal of Soluble Microbial Products. ‘Spnt’ = negative control resuspended in supernatant. ‘0.8x’ = cultures resuspended in 0.8x BG11. ‘DCMU’ = cultures resuspended in 0.8x BG11 supplemented with DCMU. ‘Dark’ = cultures resuspended in 0.8x BG11 and then incubated in the dark. ‘D→L’ = cultures resuspended in 0.8x BG11, incubated in the dark for eight hours, and then shifted to the light. ‘Sp+1.2x’ = cultures incubated in supernatant spiked with nutrient stocks to final concentration of approximately 1.2x BG11. ‘1.2x’ = cultures resuspended in fresh 1.2x BG11. Each condition was assessed with at least four biological replicates (n = 4). Error bar corresponds to one standard deviation of the sample mean.

We also investigated the role of cellular energy in aggregation. Compared to positive controls, cultures induced for aggregation by shift to 0.8x BG11 remained suspended when incubated in the dark (5.6±6.0 % aggregation, p-value < 0.01), or when 5 μmol DCMU (3-(3,4-dichlorophenyl)-1,1-dimethylurea)was added to 0.8x BG11 cultures incubated in the light (0.9±1.6 %, p-value < 0.01). Dark conditions and DCMU both prevent photoautotrophic growth [75–77]. When cultures in 0.8x BG11 were shifted to illuminated conditions after being incubated eight hours in the dark, they eventually aggregated to the same degree as without dark incubation (46.7±9.8 %). We conclude that the aggregation phenotype requires cellular energy production; i.e. it is not caused solely by change in chemical or physical environment, such as with addition of cationic coagulants for algae dewatering [78, 79], or change in ionic strength of media directly affecting the hydrophobicity and adhesiveness of cells, as described in XDLVO theory [80].

### 2.5 Soluble Microbial Products Do Not Influence Aggregation

During our aggregation assay, conditioned supernatant was exchanged for fresh BG11. In this step, the extracellular environment was modified by removal of soluble microbial products (SMP) [81]. SMP of different bacteria include the proposed secreted inhibitors and enhancers of biofilm formation by *S. elongatus*) [63], and RPS (released polysaccharides) of *E. coli*, which could influence aggregation. The pH, salinity, and osmolarity of the culture were also altered, and could be signals for aggregation, such as by inducing a stress response [68].

We therefore tested aggregation by altering nutrient concentration without removing SMP, which was accomplished by spiking 100 mL cultures resuspended in supernatant with microliter volumes of concentrated BG11 stock solutions, to approximately 1.2x BG11 final concentration (assuming supernatants of mid-log cultures are approximately 1.0x BG11). As a control, we also tested aggregation when nutrient concentration is increased by shift to fresh BG11 at 1.2x concentration. As shown in Figure 4, we found no significant difference in aggregation in 1.2x BG11 in supernatant, compared to that in fresh 1.2x BG11. Furthermore, we show that an increase in nutrient concentration is sufficient to induce aggregation to the same degree as a decrease in nutrient concentration, regardless of presence of SMP (1-factor ANOVA). We conclude that removal of SMP had no effect on cell-cell binding under the conditions tested.

We show in Figure 5 and Supplemental Material Video 1 the phenotype of a simple and rapid flocculation assay, where cell-cell binding results in larger, buoyant flocs rather than smaller sinking aggregates. We note that the IUPAC (International Union of Pure and Applied Chemistry) does not distinguish between flocculation and aggregation, both of which refer to the formation of multi-particle clusters due to destabilization of a colloid suspension [82]. We use these different terms here as a convenient way to distinguish between two different assays. It is likely that the data collected from these two assays represent variations in degree of a single phenotype, rather than two different phenotypes.

**Fig 5.**
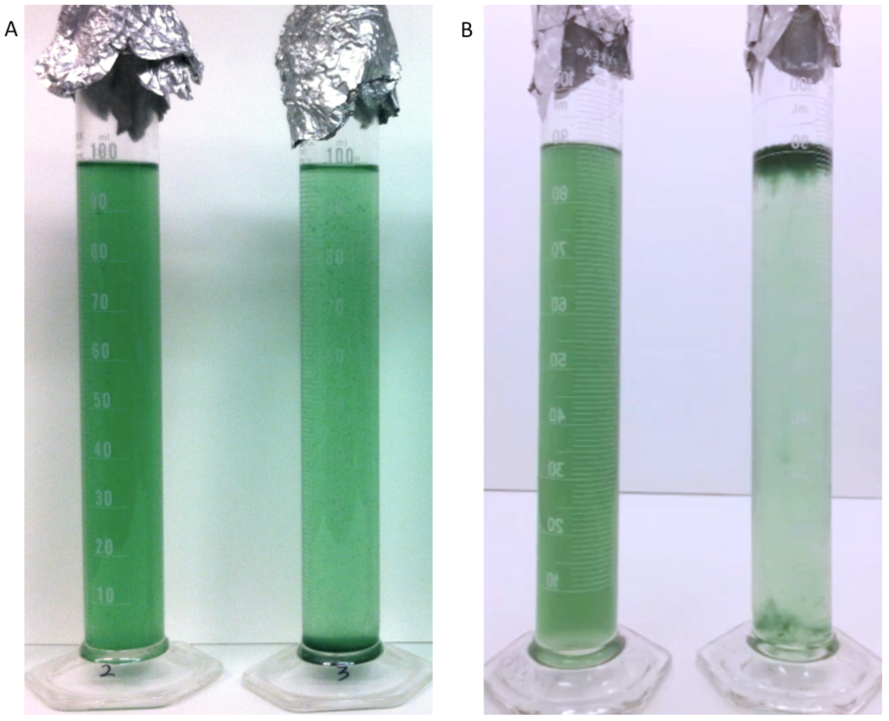
*Synechocystis* flocculation is a rapid, robust phenotype. Fig. 5A: Unaggregated WT cells in supernatant (left) and aggregated cells in 1.2x BG11 (right). Fig. 5B: Unflocculated cells in supernatants (left). Flocculation was induced by transferring culture to room-temperature autoclaved supernatants spiked with BG11 stocks to a final approximate concentration of 1.2x BG11 (right). Cell-cell binding was much faster and resulted in higher percentage of total bound biomass in WT cultures compared to the aggregation assay (representative image shown).

The flocculation assay affects a higher percent of total biomass and is a more rapid phenotype than the aggregation assay, with a duration of approximately two hours from induction to completed flocculation, compared to the aggregation phenotype which can take up to eight hours (time-course data not shown; see Supplementary Video 1 for flocculation time-lapse video). This assay may be invaluable to characterization of buoyant flocs, which are the cause of cyanobacterial surface blooms, or water blooms (blooms). Blooms and buoyant flocs have been specifically and extensively studied due to their role in harmful algal blooms (HABs), which cause malodorous and/or toxic effects with negative impact on water recreation and ecology (reviewed in (95-97)). Some phototrophs use gas vesicles to regulate their buoyancy; *Synechocystis* does not encode genes known for gas vacuole formation [83]. One study described buoyant floc formation by *Synechocystis* when 5 mmol CaCl_2_ (20x levels of 1xBG11) was added, but not BG11 up to 50x. [84] Our *Synechocystis* flocculation assay uses a nutrient strength of 1.2x BG11, resuspended in autoclaved room-temperature supernatants.

### 2.6 Type IV Pili But Not S-layer are Required for Aggregation

Figure 6 shows aggregation phenotypes between WT and mutant strains in 0.8x BG11. Aggregation by the S-layer mutant (SD523, 58.4±19.8 %) was not significantly different from WT (60.9±8.2 %). This result is not consistent with our data from the modified crystal violet assay, which showed that the S-layer mutant was inhibited for biofilm formation compared to WT (Fig. 3). As the S-layer mutant was deficient in biofilm formation but not aggregation, it is possible that S-layer is important for initial attachment, and has greater influence cell-glass binding compared to cell-cell binding. Previous studies have shown that increasing substrate surface roughness decreases the surface interaction energy, facilitating binding [85, 86]. Similarly, it may be that *Synechocystis* binding to smooth glass coverslips during the modified crystal violet assay is less favorable compared to binding to a rough cell surface during the aggregation assay. Additionally, there is precedence for influence of the pH and ionic strength of the growth medium on the binding phenotypes of S-layer mutants, including studies of *Lactobacillus, Clostridium*, and *Geobacillus*, three Gram-positive genera [80, 87]. If this variation in hydrophobicity also affects *Synechocystis* S-layer mutants, it may play a larger role in binding to glass than binding to other cells.

**Fig 6.**
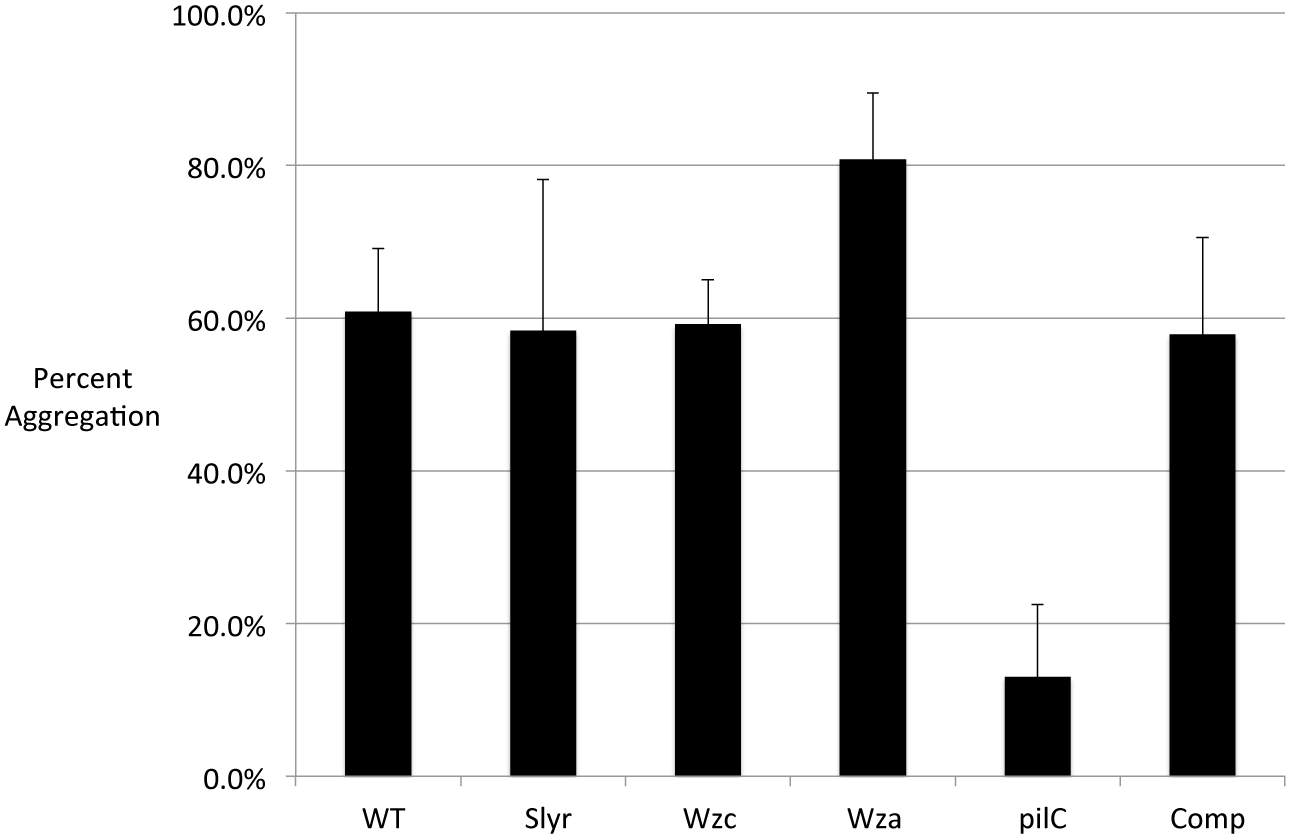
Cell Surface Structures Differentially Modulate *Synechocystis* Aggregation. WT and mutant strains analyzed with aggregation assay (shift to 0.8x BG11). ‘WT’ = SD100; ‘Slyr’ = SD523; ‘Wzc’ = SD546; ‘Wza’ = SD517; ‘pilC’ = SD519; ‘Comp’ = SD577. Error bar corresponds to one standard deviation from the sample mean.

The *pilC* deletion mutants (SD519) were severely deficient in aggregation (13.0±9.5 %, compared to WT 60.9±8.2 %, p-value = 0.01). This is consistent with the *pilC* mutants also being deficient in biofilm formation. Introducing the *pilC* gene and putative promoter region back into the *pilC* mutants via expression plasmid restored WT levels of aggregation (SD577, 57.9±12.7 %). *Synechocystis* pili extend several microns beyond the cell surface in all directions [27, 59]. The role of pili in WT aggregation thus may partly be due to increasing the effective volume occupied by each piliated cell, thereby promoting cell-cell contact. The filamentous shape of pili also facilitates binding by reducing the effective radius at the point of surface contact with pili tips, as predicted by DLVO theory; this is also consistent with pili playing a larger role than S-layer in aggregation (Figure 5). Since we showed that SMP do not influence aggregation (Figure 4), our data preclude PilC-mediated secretion of released factors analogous to biofilm enhancers and inhibitors secreted by *S. elongatus* [62–64].

### 2.7 Mutants Lacking Wza-dependent EPS Have a Super-binding Phenotype

Mutants lacking *wzc* (SD546) had WT levels of aggregation (59.2 ±5.8 %), whereas mutants lacking *wza* had significantly higher levels of aggregation (80.8±8.7 % vs 60.9±8.2 %, p-value = 0.01). This is consistent with previously reported results of EPS affecting cell-cell repulsion [29, 38]. In one study, a 3-week standing flask assay of cultures in unaltered supernatants showed that mutants lacking Wza fell out of suspension (sedimented without aggregation) more readily than WT or Wzc mutants. Wza mutants were also shown to have less EPS, and a smaller zeta-potential (approximately −21 mV) than WT (approximately −35mv) [38]. Zeta-potential is a proxy for quantifying the electrostatic charge of cell surfaces [88]. The authors of the study concluded that Wza-dependent EPS production promotes dispersal of planktonic WT cells via electrostatic repulsion, consistent with the super-binding phenotype of the *wza* deletion mutants reported here. This conclusion is also consistent with a third study reporting constitutive aggregation and binding to glass of *Synechocystis* mutants lacking EPS [28]. The mutations in *wzt/kpsT* and *wzm/kpsM* are predicted to function as ABC-transporters in the same pathway as *wza* and *wzc*.

### 2.8 No Differential Expression of Cellulose or Outer Membrane Proteins Detected During Aggregation

As with biofilm formation, Type IV pili are necessary but not sufficient for aggregation by WT *Synechocystis*: an additional factor, such as induced by change in concentration of BG11, is required for aggregation to occur. We hypothesized that WT *Synechocystis* was producing adhesive molecules on its surface in response to changes in growth medium, causing aggregation. To detect these putative adhesive molecules, we isolated outer membrane protein fractions of WT and mutant strains under treatment and control aggregation conditions. Samples were split before loading on gels to include both boiled and unboiled preparations of proteins, in order to detect any heat labile proteins. As shown in Supplementary Material Figure S1, we did not detect any putative proteins that were differentially expressed in the treatment (aggregated) cultures vs negative control, as determined by altered band patterns of Coomassie stained gels analyzed by SDS-PAGE. This result may indicate that aggregation is not mediated by synthesis of an outer membrane protein adhesin, but mass spectrometry would be more definitive.

Aggregated cells of *Thermosynechococcus vulcanus* RKN were visibly dispersed by treatment with cellulase enzyme, consistent with the role of cellulose in causing aggregation in this species [40]. We used the published cellulase assay to test for role of cellulose in *Synechocystis* aggregation. Compared to a negative control, we saw no significant change in degree of aggregation in cellulase-treated cultures (data not shown). As shown in Supplementary Material Figure S2, we also tested for presence of cellulose in the purified extracellular matrices of aggregated and control cultures using the glucose oxidase assay, which quantifies amount of glucose released by digestion with cellulase enzyme, as shown previously with *T. vulcanus* RKN samples [40]. While our results do indicate that cellulase-liberated glucose was present in our samples compared to negative and positive controls (p-values < 0.03), these glucose equivalent levels showed no significant variation between aggregated and unaggregated samples. A mutational analysis of the biofilm and aggregation phenotypes of *Synechocystis* mutant lacking the gene *sll1377*, which contains the putative cellulose synthase domain, would be more definitive.

Based on this and previous studies, our current understanding is that *Synechocystis* aggregation is bioenergy-dependent, is induced directly or indirectly by changes in nutrient concentrations but not SMP removal, in a process that requires Type IV pili but not cellulose, and is inhibited by Wza-dependent cell-bound EPS. Additionally, we show in Supplementary Figure S3 that mutants lacking S-layer and Wza phenocopied WT for gliding motility and phototaxis. This indicates that the influence of these mutations on biofilm formation and aggregation is not due to epistatic effects on motility pathways that may be indirectly required for cell binding. Additionally, our WT Kazusa strain (SD100, see Table 1) with amotile Type IV pili is competent for aggregation and biofilm formation, indicating motility is not required for cell binding in *Synechocystis*.

Although the precise molecular mechanisms of *Synechocystis* cell binding remain to be determined, our data are consistent with the relative importance of these factors as reported in other bacteria: cellular energy/growth was essential for any aggregation to occur, whereas the smaller effects of S-layer and Wza-dependent EPS on aggregation and biofilm formation suggest their role is due to electrostatic and/or hydrophobic contributions to the cell surface. Likewise, our data indicate that pili have a larger impact than S-layer or Wza-dependent EPS, which could be attributed to their additional roles in increasing effective culture density, and / or reducing the effective radius of the point of surface contact, which is predicted by DVLO theory to increase binding.

## 3.0 Conclusion

Our results further the development of *Synechocystis* as model organism for studies of axenic phototrophic biofilm formation and aggregation. We report convenient new axenic assays relevant to ecological and biotechnological studies of cell binding under controlled laboratory conditions, namely a modified crystal violet assay for biofilm formation, in addition to aggregation and flocculation assays. These new assays enable much more rapid analysis (<72 hours vs weeks) of WT *Synechocystis* cell binding phenotypes compared to those published previously. This is due to using changes in nutrient concentration to induce binding of exponentially growing cultures, rather than growing cultures in blue light [50] or light-activated heterotrophic growth (LAHG) [49], which necessitates using slow-growing cultures.

We demonstrate the utility of these assays in performing mutational analysis to identify cell surface structures influencing cell-cell binding, namely Type IV pili, Wza-dependent exopolysaccharide, and S-layer. These findings include the report of a non-biofouling strain of *Synechocystis*, the *pilC* deletion mutant SD519, which would be an advantageous genotype for feedstock strains cultivated in planktonic PBRs or open ponds. Additionally, we used these assays to determine that change in nutrient concentration of *Synechocystis* cultures is an immediately useful environmental signal for rapid, economical harvest of 60 % biomass of WT *Synechocystis* cells (or 80 % of *wza* mutant cells) within eight hours. These assays also comprise new tools for performing studies on molecular biology of axenic cell binding by a phototrophic bacterium, which historically have been primarily conducted with axenic heterotrophic bacteria.

Overall, our data and previously published studies are consistent with a model for WT *Synechocystis* cell-cell interactions regulated by two different mechanisms depending on growth conditions. Under optimal growth conditions, the negatively charged Wza-dependent EPS keeps cells distributed in a stable colloid suspension by electrostatic repulsion as predicted by XDLVO theory; this could benefit the cell by limiting self-shading that would otherwise be caused by sedimentation [38] or transient contact in cell suspensions [89]. Under altered nutrient conditions, blue light, or LAHG, this cell-cell repulsion is overcome through an unknown mechanism that, based on studies in other bacteria, likely includes synthesis and export of adhesive molecules to the cell surface.

In oligotrophic natural environments such as lakes and pelagic zones of open oceans, cyanobacterial cell density is much lower than that typically used for lab cultures [90], reducing the number and size of aggregates detected in these natural environments [72]. However, migration of cyanobacteria through the water column is a normal part of their seasonal adaptation, forming blooms on lake surfaces in spring, and benthic layers in the winter [90–92]. Aggregation under nutrient-limited conditions contribute benefits to phototrophs (reviewed in [93]), including relocation of cells to nearby micro-niches that may not be as nutrient-limited. Cyanobacterial aggregates (particulate organic carbon) also have important ecological implications, as they have recently been identified as important contributors to carbon flux to lower ocean depths, which has a major impact on oceanic food webs [94, 95]. Global warming may disrupt these natural cycles in a number of ways, including increased temperatures and ocean acidification; overall, climate change is predicted to increase the growth of cyanobacterial blooms, including those species known to be toxic [96, 97]. Additional studies will be helpful in developing strategies to mitigate these negative effects on ocean food webs, and in optimizing cyanobacteria for production of sustainable food, fuel, and other valuable commodities [98, 99].

## 4.0 Materials and methods

### 4.1 Culture Growth Conditions

*Synechocystis* cultures were grown in 100 mL of BG11 medium [100] in 500 mL Erlenmeyer flasks until they reached to mid-log (OD_730_ 0.6 to 0.8). Cultures were illuminated continuously with 50 μmol/(m^2^ s) photons (PAR) and mixed at 120 rpm on a platform shaker. Cultures were bubbled with a supply of sterile, humidified air at 0.8 mm/min as measured by a flow meter (Cole Palmer). Air was sterilized by passing through a 0.22 micron filter, and pre–humidified by bubbling through a side–arm flask of sterile dH_2_O. The flasks were incubated in growth chambers maintained at constant 30% humidity. The cultures were diluted to an OD_730_ of approximately 0.15 and allowed to double at least twice before assessing cell–binding phenotypes. Medium was supplemented with 50 μg/mL kanamycin sulfate for growth of mutant strains carrying a kanamycin resistance marker. Strain SD577, carrying plasmid pΨ552, was propagated with media containing 30 μg/mL each of streptomycin and spectinomycin.

### 4.2 Modified Crystal Violet Assay

A standard crystal violet assay was adapted to *Synechocystis* from methods described previously (‘microtiter dish biofilm formation assay’, [66], ‘plastic binding assay’ [67]. Cultures were diluted in 100 mL of BG11 at a starting OD_730_ of approximately 0.15 and grown again to log phase as follows: for negative control cultures (uninduced), growth conditions were as described above; for treatment cultures (induced), the side-arm flask for humidification was removed, resulting in evaporation of culture flask to about 84 mL volume over 24 hours, equivalent to a nutrient strength of approximately 1.20x BG11. Therefore, returning this culture to 1.0x BG11 at the start of a biofilm assay (described below) introduces a shift from higher to lower nutrient condition.

Cells from control and treatment cultures were harvested by centrifugation at 6,000xg for 5 minutes and resuspended in 1.0x BG11 to an OD_730_ of 0.05. Three mL of culture was added to each well of a 12 well plate (Corning Costar, Fisher Scientific catalog number 07–200–82) that contained a 22–mm thick glass coverslip as biofilm substratum (Fisher Scientific catalog number 12–540–B). Glass coverslips were trimmed previously to fit vertically into the plate wells using a diamond scribe (Ted Pella, Inc.). Plates with inserted coverslips were placed in cross–linker (Spectroline Spectrolinker XL–1500). Plate lids were removed and also placed face up in cross–linker. Materials were sterilized by UV radiation (254 nm) for 400 seconds at 1,500 1500 μW/cm^2^ (600 mJ/cm^2^). After inoculation of wells, tissue plate edges were sealed to plate lids with Parafilm and cellophane tape to minimize evaporation. Sealed plates were incubated for 72 hours on platform shaker at 72 rpm under 32 μmol/m^2^/ sec photons (PAR), in a chamber with 30% humidity. 1.0x BG11 medium without inoculum was used as a blank.

Coverslips were removed and rinsed 10 seconds per side with strong stream of BG11 from a squeeze bottle, and excess solution was wicked off by standing the coverslip edgewise on absorbent paper for 5 seconds. Coverslips were then stained by inserting in wells containing 4 mL of 0.01% aqueous crystal violet solution (w vol) for five minutes in a separate tissue culture plate. Unbound stain was rinsed and wicked away, as above. Coverslips were dried in ambient air overnight in the dark and used for qualitative assessment (imaging of macroscopic staining patterns). The final culture OD_730_ of each well was also measured to correlate planktonic culture growth with biofilm growth. For quantifying biofilms, dried coverslips were placed in small weigh boats and crystal violet stain was dissolved in 1 mL of DMSO with platform shaking for 20 minutes in the dark or until coverslip stain was removed (up to 45 minutes). Crystal violet absorbance was measured at 600 nm as a proxy for the amount of cellular material bound to coverslips.

### 4.3 Confocal Microscopy Imaging of Biofilms

Biofilms were grown on glass coverslips and rinsed (without staining) as described in the modified crystal violet assay (above) and placed on ice to cool. 100 μL of 1.6% low-melt agarose in sterile isotonic solution (1% NaCl w vol in dH_2_O) was immediately applied to the biofilm facing up on the chilled coverslip. Coverslips with agarose facing up were then placed in a small petri dish and attached to dish with dental wax and immersed in isotonic solution. Biofilms were imaged using a Leica TCS SP5 II with 10x or 63x DIC dipping lens. Sample fluorescence was excited by argon laser and collected through Texas Red filter. Biofilm heights were calculated manually by measuring samples of interest using calibrated Z-axis.

### 4.4 Plasmid and Strain Construction

DNA manipulation was carried out using standard procedures [101]. Suicide plasmids for replacing *Synechocystis* genes with a KmR*sacB* cassette were constructed by four-part ligation into commercial vector pGEM 3Z (Promega), a pUC18 derivative. The KmR*sacB* cassette from pPSBA2ks [69] contains markers for kanamycin resistance and sucrose sensitivity. PCR fragments of approximately 500 base pairs (bp) located upstream and downstream of each gene target were amplified from the *Synechocystis* genome using primers from Supplementary Material Table S1 to target locations of double homologous recombination flanking the gene of interest for each suicide vector. Flanking regions, pGEM 3Z, and the KmR*sacB* cassette were stitched together by restriction digest and ligation as follows. Briefly, NheI– and EagI–digested restriction sites were generated between the two flanking sequences to accommodate the digested KmR*sacB* fragment, and BamHI and SphI (New England Biolabs) sites allowed insertion of these three fragments into the pGEM 3Z multi–cloning site. For example, we created the plasmid p*ψ*541 (for replacing *pilC* with KmR*sacB*) by ligating digested PCR products amplified from upstream and downstream regions of *pilC* with digested KmR*sacB* and pGEM 3Z. (For replacing *wzc*, BamHI and XbaI were used to insert the KmR*sacB* cassette, and SacI and SphI were used to insert the fragments to create pΨ546). Ligation reactions were transformed into competent *E. coli* (5–alpha, New England Biolabs) and transformants were screened by antibiotic selection. Clones were identified by DNA sequencing.

Design of KmR*sacB* suicide plasmids incorporated genomic context of each gene to avoid introducing polar effects of neighboring genes as follows. For *sll1581*, the last 100 bp were left intact in order not to delete putative upstream promoter–containing region of neighboring *sll1582*. Similarly, the putative promoter regions of *sll1581* and *ssr2843* overlap, therefore this region was not included in the replacement region in order to preserve native expression of *ssr2843* in *sll1581* deletion mutants. For the remaining mutant strains, gene and upstream region (predicted promoter) were targeted for KmR*sacB* replacement.

### 4.5 Transformation of Naturally Competent or Electrocompetent *Synechocystis* Cultures

Mutants of *Synechocystis* were generated as previously described [102]. Briefly, cells from log-phase culture were harvested as above and resuspended to 200 μL volume equivalent of OD_730_ = 2.5. Four μg of suicide vector was added to *Synechocystis* cells and incubated for six hours in BG11 without antibiotic, with intermittent shaking. The transformation reaction was plated onto a Nuclepore Track-etch membrane (Whatman) on a BG11 agar plate. These were then incubated for 24-72 hrs at 30°C with 50 μmol/(m^2^ s) photons PAR, until a green lawn appeared. Following this incubation, membranes were transferred to a BG11 plate containing 50 μg/mL kanamycin sulfate and incubated for approximately 2 weeks until lawn disappeared and then single colonies grew. To ensure complete segregation of the transformants, colonies were re-propagated three to five times on BG11 agar plates supplemented with 50 μg/mL kanamycin sulfate. Kanamycin–resistant colonies were screened via PCR with primers internal to genes targeted for deletion to determine complete segregation of chromosomes, indicating loss of the WT gene sequence.

Since natural competence in *Synechocystis* requires Type IV pili [59], we prepared electro–competent cultures of our apiliate *pilC* mutants in order to transform them with plasmid pΨ552 for expressing PilC, as described previously [103, 104]. We harvested cells from 50 mL of log-phase cultures as described above. Cells were resuspended in 500 μL of sterile 10% glycerol solution. 60 μL samples of cells were mixed with up to 10 μg purified plasmid (300-3000 ng/μL DNA in dH_2_O). Cells and DNA were added to 0.1 cm electroporation cuvettes and then pulsed with 12 kV/cm, 25 μF, 400 Ω setting. Cells were resuspended in 900 μL BG11, transfered to test tubes with 2 mL additional BG11, and incubated as described previously until OD_730_ doubled. To select for transformants, cells were harvested by centrifugation and resuspended in 500 μL supernatant for plating a range of volumes on selective medium (BG11 supplemented with 30 μg/mL each of streptomycin and spectinomycin).

### 4.6 Isolation and Analysis of Outer Membrane Protein Fractions

Cells were lysed and fractionated using standard methods [105]. 15 mL volumes of cultures were harvested by centrifugation at 6,000xg for 5 minutes; cells were stored at 80°C. Cells were resuspended in 1.2 mL of 50 mM ammonium bicarbonate buffer solution with HALT protease inhibitor (ThermoFisher) on ice. 600 μL of sample were added to 2 mL cryovials with 400 μL of 0.1 mm zirconium beads. Cells were lysed by bead–beating (Mini BeadBeater, BioSpec) for 7 cycles at maximum speed (one cycle is 30 seconds beating followed by 2 minute incubation on ice). Whole–cell lysates were fractionated using differential centrifugation as follows. Lysates were transferred to new tubes. Unlysed cells were harvested at 1,600xg for 5 minutes. Supernatants were transferred and total membrane fraction was harvested at 16,000xg for 1 hour. Total membrane pellet was resuspended in 500 μL of 20 mM Tris pH 8 and 500 μL 0.8% Sarkosyl on ice, and incubated at 4°C with inversion for 90 minutes. Outer membrane fraction was pelleted at 16,000xg for 8 h to 12 h at 4°C. Supernatant was removed, and final pellet of enriched outer membrane fraction was resuspended in 50 μL of 20 mM tris buffer pH 8. Samples were quantified via BCA assay (Sigma Aldrich). Volumes equivalent to equal protein (20 μg per well) were added to SDS loading buffer, heated (or not heated) at 100°C for 10 minutes, and loaded onto gradient acrylamide gels for PAGE. Protein bands were visualized via Coomassie stain [101].

### 4.7 Aggregation assay

Cells from 100 mL of mid-log culture (OD_730_ of 0.6-0.8) were harvested by centrifugation as described above, resuspended in either supernatant (negative control) or 0.8x BG11 (treatment condition), and decanted into 100 mL glass graduated cylinders. The starting OD_730_ was measured, and the standing cultures were incubated at 30°C with illumination as described above. The final OD_730_ after eight hours was measured by sampling 1 mL of culture from the 50 mL mark on the cylinder. Aggregation was reported as normalized percentage change in OD as follows: [(Final OD – Starting OD) / Starting OD] x100. Negative % aggregation indicated that the culture density increased over time due to cell growth, while minimal aggregation occurred.

### 4.8 Flocculation Assay

*Synechocystis* cultures were grown to midlog phase (OD_730_ between 0.6 to 0.8), and cells were harvested as described above. Supernatants were decanted into large flasks (5 liters or more); capped with foil and placed in secondary containment pans. Supernatants were autoclaved for five minutes on gravity cycle with no drying (total time of autoclave cycle should be no more than about 15 minutes.) Supernatants were cooled to 30°C with ice bath using a digital thermometer to monitor temperature.

BG11 stock solutions were added to cooled supernatants to increase medium strength to a final approximate concentration of 1.2x BG11, assuming supernatant contributes approximately 1.0x BG11. Harvested cells were resuspended in the prepared supernatants. Cultures were then decanted into graduated cylinders, and incubated and measured as described above.

### 4.9 Cellulase Treatment of Aggregated Cultures

Cellulase digestion of aggregates was performed as described previously [40]. Three mL of aggregated cultures were transferred to 15 mm glass test tubes, and 100x stock solutions of cellulase were prepared in dH_2_O and added to final concentrations of 0.60 U/mL cellulase (Sigma-Aldrich, product number C0615). An equal volume of water was added to a negative control culture. Cultures were mixed and incubated without shaking at 30°C in the light. After incubation, cultures were gently resuspended, and 1 mL samples were transferred to semi–micro cuvettes and allowed to settle for one hour. Dispersal of aggregates was determined by increase in absorbance at OD_730_ between treated and control condition.

### 4.10 Isolation of Extracellular Matrix for Biochemical Characterization

Extracellular matrix (ECM) of cells, including S–layer and exopolysaccharide, were removed and purified by a combination of mechanical and chemical separation from intact cells as described previously [28, 29, 106]. 100 mL cultures of OD_730_ approximately 0.6-0.8 were centrifuged for 20 minutes at 6,000xg. Cells are resuspended in 10 mL of supernatant with 60 μL of formaldehyde and incubated at 4°C for one hour. 4 mL of 1 N NaOH were added, mixed gently and incubated again at 4°C for three hours to disrupt ionic bonding between the ECM and the cell. Cells were centrifuged for 20 minutes at 20,000xg to physically separate ECM from cells. Supernatants were passed through 0.2 micron filters to remove trace cells. A negative control of BG11 medium was also sterile–filtered to detect background levels of cellulose from the filtration membrane, and*\*or dialysis membrane (see below). Samples and negative controls were lyophilized, resuspended in 2 mL dH_2_O, and dialyzed into 3×4 L dH_2_O using 4000 MWCO reconstituted cellulose membranes (Tube–O–Dialyzer, G Biosciences, catalog number 786–616). Samples were then lyophilized a second time, resuspended in 100 μL dH_2_O, and stored at 20 μL aliquots at *−*80°C until analysis.

### 4.11 Digestion and Quantitation of Cellulose from ECM using Glucose Oxidase Assay

Cellulose from ECM was digested and quantified as described previously [40], with modifications. ECM samples and cellulose positive control (Sigma-Aldrich product number C8002) were digested with cellulase enzyme (Sigma-Aldrich, product number C0615) to liberate glucose, which was detected by a glucose oxidase assay. Reactions were prepared in 100 μL with a final concentration of 4% w\v cellulose polymer (positive control) or 0.89x ECM. Cellulase enzyme was prepared in 40 mM sodium acetate buffer, and added to reactions for a final cellulase concentration of 0.6 U/mL. Reactions were digested at 37°C for 72 hours. Digested samples are centrifuged at 13,200 xg for one minute and supernatants were transferred to new tubes and stored at 80°C.

To measure glucose released by digestion with cellulase enzyme, the BioAssay Systems EnzyChrom Glucose Oxidase Assay Kit (EGOX–100) was used according to manufacturer’s instructions as described previously for cyanobacterial samples [40], with modification. Working reagent was prepared to include glucose oxidase enzyme, and exclude addition of 2 M D-glucose solution. Glucose Oxidase Enzyme solution (Sigma G0543-10KU) was diluted in dH_2_O to final reaction concentration of 8.3 U/mL of glucose oxidase. Working reagent was combined with 20 μL of 1x digested ECM or 1:200 dilution of digested cellulose polymer positive control in 96-well plate. Plates were covered with lids and vortexed gently, centrifuged briefly, and then incubated at room temperature for 20 minutes. Fluorescence was measured at 530ex 585em with a SpectraMax M5 plate reader (Molecular Devices) using Softmax Pro software version 6.2.2.

### 4.12 Phototaxis and Motility Assay

The phototaxis assay was adapted from Bhaya et al. [61]. Log–phase cultures were diluted to OD_730_ of about 0.25, and 10 μL volumes were spotted on swarm agar (BG11 medium prepared with 0.5% Difco BactoAgar) in grid-lined square petri dishes, such that inocula were oriented directly over grid lines. Plates were sealed in parafilm and incubated at 30°C and 30% ambient humidity under directional illumination (a dark box with a 30 μmol/(m^2^ s) photons PAR light source at one end). Strains were graded as motile and phototactic using qualitative assessment of growth having blurred edges, and having elongated away from grid line, as compared to negative control strain, which grows in a disc with crisp edges centered on top of grid line (See Supplemental Material Figure S3).

## 5.0 Supplementary Material

**Video S1. Time-Lapse Video of Flocculation Assay**

Left, treatment condition: WT *Synechocystis* cultures suspended in autoclaved room-temperature supernatant spiked with BG11 stock solutions to final concentration of 1.2x BG11. Right, negative control condition: WT *Synechocystis* cells were resuspended in unautoclaved supernatant with no added BG11 stocks. One hour time lapse, starting at time = 1 hour and ending at time = 2 hours.

**Fig. S1.**
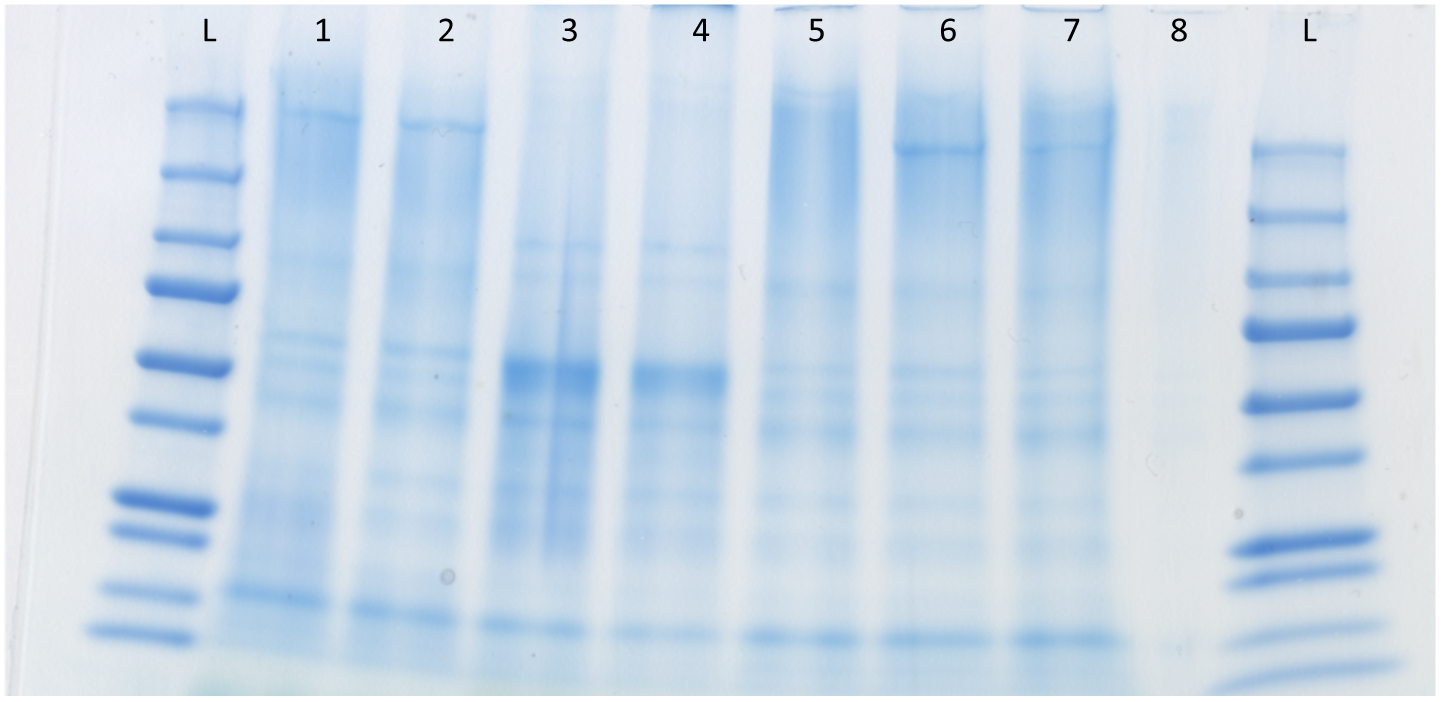
Coomassie Stain of Outer Membrane Proteins Analyzed by SDS-PAGE. Induced = incubated in 0.8x BG11; Uninduced = incubated in supernatant, as in aggregation assay. L = Ladder, 1 = WT Uninduced; 2 = WT Induced; 3 = WT Uninduced (OMP not boiled); 4 = WT Induced (OMP not boiled); 5 = S-layer mutant (SD523) Induced; 6 = PilC mutant (SD519) Induced, 7 = PilC Comp (SD577) Induced, 8 = Empty Lane. Ladder bands range in size from 10 to 250 kDa.

**Fig. S2.**
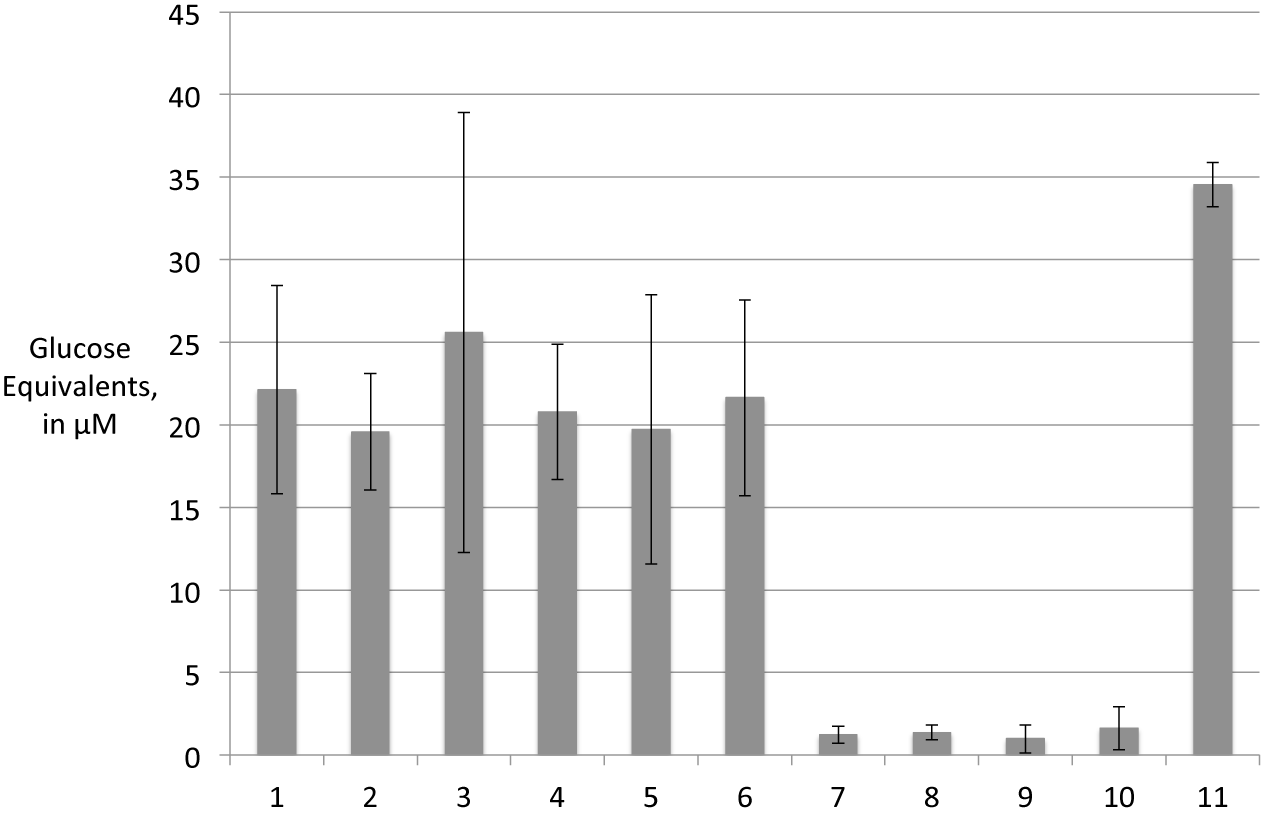
Glucose Oxidase Assay of Extracellular Matrices Digested with Cellulase. Extracellular matrices (ECMs) of WT and mutant cultures from aggregation assays in Fig. 4 and 5 were isolated and digested with cellulase enzyme to liberate glucose. Glucose was detected indirectly by fluorescence measurement in glucose oxidase assay and compared to positive controls (digestion of pure cellulose) and negative controls (no cellulase enzyme; no ECM), relative to pure glucose standard solutions. Induced = 0.8xBG11; Uninduced = supernatant, as described in aggregation assay. ECMs: 1 = WT (SD100) 0.8x BG11 (Induced); 2 = WT supernatant (Uninduced); 3 = PilC (SD519) (Induced); 4 = pilC Comp (SD77) (Induced); 5 = Wza (SD517) (Induced); 6 S-layer (SD523) (Induced). Controls: 7 = Sterile-filtered and dialyzed BG11; 8 = Water blank; 9 = Cellulase enzyme -, cellulose +; 10 = Cellulase enzyme +, Cellulose -; 11 = Cellulase enzyme +, Cellulose + (1:2000 digest dilution).

**Fig. S3.**
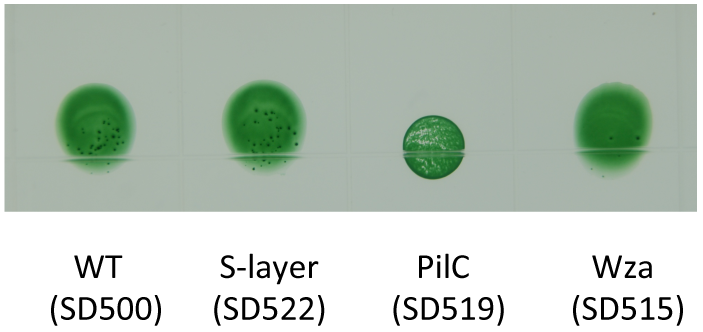
Motility and Phototaxis Assay of WT and Mutant *Synechocystis* strains.

**Table S1.**
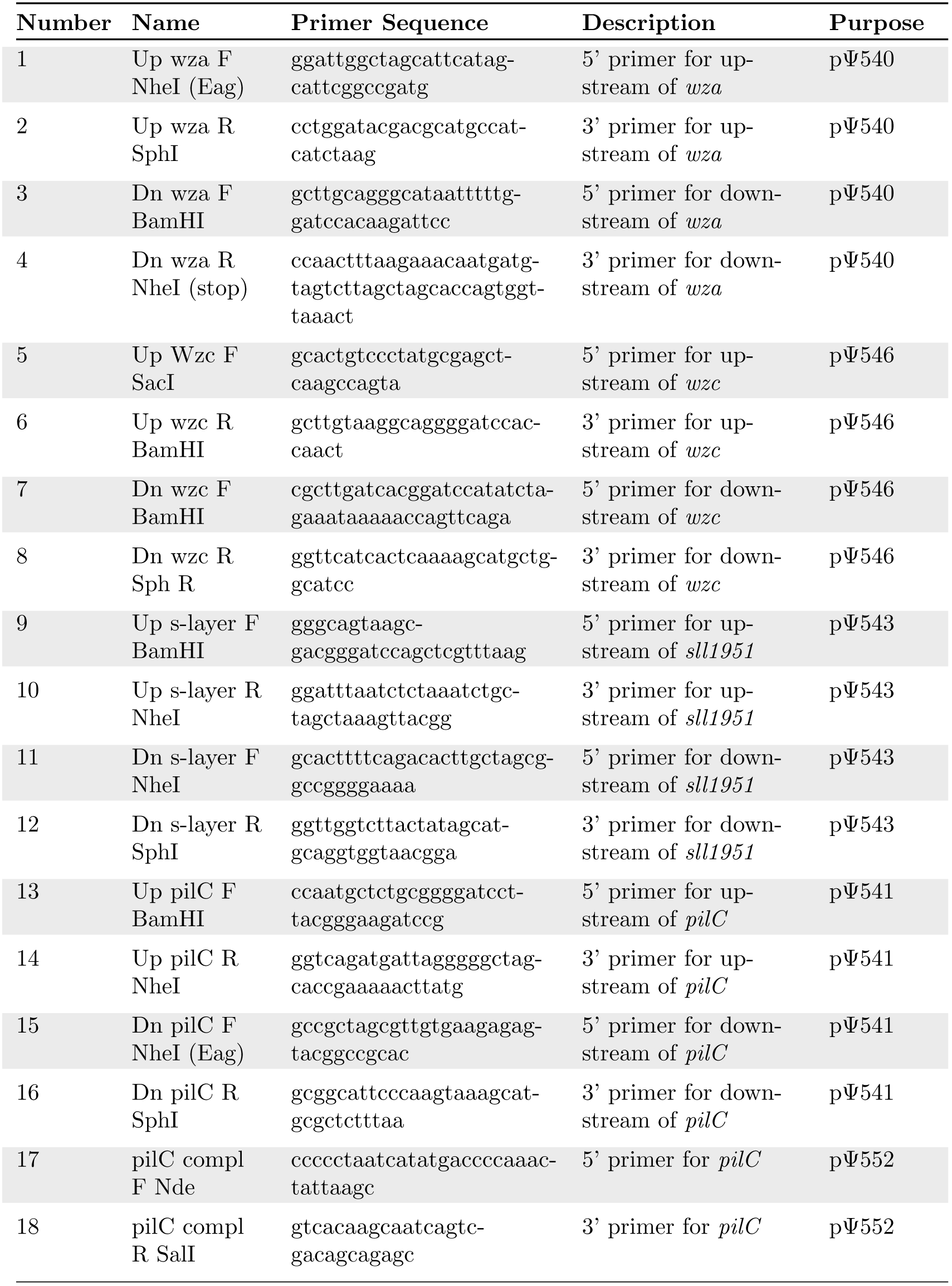
Primers Used in This Study.

## Acknowledgments

We wish to gratefully acknowledge the expertise and assistance in confocal microscopy of Dr. Debra Baluch, at the Keck Imaging Facility in the School of Life Sciences at Arizona State University, Tempe. We also are indebted to Dr. René Daer for essential assistance and invaluable suggestions for manuscript preparation. Dr. Penny Gwynne, Dr. Wei Kong, and Dr. Shelley Long in The Biodesign Institute at Arizona State University each generously shared access to essential equipment for lyophilization and other sample preparation. We are grateful to Dr. Willem Vermaas and Dr. Soo-Young Wanda for sharing essential strains and plasmids.

